# KLF4 in smooth muscle cell-derived progenitor cells is essential for angiotensin II-induced cardiac inflammation and fibrosis

**DOI:** 10.1101/2024.06.04.597485

**Authors:** Sizhao Lu, Austin J Jolly, Allison M Dubner, Keith A Strand, Marie F Mutryn, Tyler Hinthorn, Tysen Noble, Raphael A Nemenoff, Karen S Moulton, Mark W Majesky, Mary CM Weiser-Evans

**Affiliations:** Department of Medicine, Division of Renal Diseases and Hypertension, University of Colorado Anschutz Medical Campus, Aurora, CO, USA; Medical Scientist Training Program, University of Colorado School of Medicine, Anschutz Medical Campus, Aurora, CO, USA; Biomedical Sciences and Biotechnology MS program, University of Colorado Graduate School, Anschutz Medical Campus, Aurora, CO, USA; Department of Medicine, Division of Cardiology, University of Colorado Anschutz Medical Campus, Aurora, CO, USA; School of Medicine, Consortium for Fibrosis Research and Translation, University of Colorado Anschutz Medical Campus, Aurora, CO, USA; Cardiovascular Pulmonary Research Program, University of Colorado Anschutz Medical Campus, Aurora, CO, USA; Center for Developmental Biology & Regenerative Medicine, Seattle Children’s Research Institute, Seattle, WA 98101; Departments of Pediatrics, Laboratory Medicine & and Pathology, University of Washington, Seattle, WA, 98195

**Keywords:** cardiac fibrosis, smooth muscle-derived stem cell, reprogramming, scRNA-Seq periostin

## Abstract

Cardiac fibrosis is defined by the excessive accumulation of extracellular matrix (ECM) material resulting in cardiac tissue scarring and dysfunction. While it is commonly accepted that myofibroblasts are the major contributors to ECM deposition in cardiac fibrosis, their origin remains debated. By combining lineage tracing and RNA sequencing, our group made the paradigm-shifting discovery that a subpopulation of resident vascular stem cells residing within the aortic, carotid artery, and femoral aartery adventitia (termed AdvSca1-SM cells) originate from mature vascular smooth muscle cells (SMCs) through an *in situ* reprogramming process. SMC-to-AdvSca1-SM reprogramming and AdvSca1-SM cell maintenance is dependent on induction and activity of the transcription factor, KLF4. However, the molecular mechanism whereby KLF4 regulates AdvSca1-SM phenotype remains unclear. In the current study, leveraging a highly specific AdvSca1-SM cell reporter system, single-cell RNA-sequencing (scRNA-seq), and spatial transcriptomic approaches, we demonstrate the profibrotic differentiation trajectory of coronary artery-associated AdvSca1-SM cells in the setting of Angiotensin II (AngII)-induced cardiac fibrosis. Differentiation was characterized by loss of stemness-related genes, including *Klf4*, but gain of expression of a profibrotic phenotype. Importantly, these changes were recapitulated in human cardiac hypertrophic tissue, supporting the translational significance of profibrotic transition of AdvSca1-SM-like cells in human cardiomyopathy. Surprisingly and paradoxically, AdvSca1-SM-specific genetic knockout of *Klf4* prior to AngII treatment protected against cardiac inflammation and fibrosis, indicating that *Klf4* is essential for the profibrotic response of AdvSca1-SM cells. Overall, our data reveal the contribution of AdvSca1-SM cells to myofibroblasts in the setting of AngII-induced cardiac fibrosis. KLF4 not only maintains the stemness of AdvSca1-SM cells, but also orchestrates their response to profibrotic stimuli, and may serve as a therapeutic target in cardiac fibrosis.

## INTRODUCTION

Cardiac fibrosis is one of the common end-stage pathologies that characterize most cardiovascular diseases, such as heart failure, hypertension, and coronary artery disease^1,2^. Although fibrotic tissue maintains organ integrity, excessive deposition of extracellular matrix (ECM) in cardiac tissue significantly disrupts normal function of the heart^3^. Traditionally, fibroblasts are recognized as the major contributor to cardiac fibrosis through their activation to a myofibroblast phenotype^1,2,4^. Compared to their quiescent counterparts, myofibroblasts exhibit increased ECM production, cytokine and chemokine secretion, and increased expression of muscle contractile genes including alpha Smooth Muscle Actin (αSMA). This simplified view, however, has been increasingly challenged by recent findings, especially with the employment of single-cell profiling technologies, which have revealed the innate heterogeneity of fibroblast populations^2,5,6^. Mouse cardiac fibroblasts exhibit a spectrum of transcriptomic profiles suggesting their unique functions in different fibrotic disease settings^2,7–9^. Similarly, large scale collaborative scRNA-seq profiling of human cardiac tissue revealed distinct fibroblast populations, including periostin (*POSTN*)-expressing, ECM-producing/remodeling, cytokine receptor (*OSMR* and *ILST6*)-expressing, and mesenchymal stem cell-like(*CD34*^+^) cells^10^. The heterogeneity of cardiac fibroblasts poses a major challenge in accurately quantifying and defining their function in disease settings^5^.

The vascular adventitia is the main site of pathological fibrotic changes that reduce arterial compliance^11–13^. The adventitia is a dynamic and complex layer with a variety of cell types including immune cells, fibroblasts, a microvascular network known as the vasa vasorum, and importantly, resident progenitor cells^14,15^. Resident adventitial progenitor cells express the stem cell marker Sca1^+^ (AdvSca1 cells) and play a vital role in growth, remodeling, repair, and injury of the artery wall^16–18^. Vascular medial smooth muscle cells (SMCs) are specialized cells that express high levels of SMC-specific contractile proteins, but under pathological conditions are capable of undergoing profound phenotypic and functional changes^19^. We previously demonstrated that, in a physiological process, mature SMCs migrate into the adventitia, are reprogrammed in a KLF4-dependent manner into a distinct subtype of AdvSca1 progenitor cells termed AdvSca1-SM cells, and reside in an adventitial progenitor niche; KLF4 is also essential for the maintenance of the stemness phenotype of AdvSca1-SM cells^20,21^. Further, we validated that a *Gli1*-promoter-driven inducible Cre mouse model is a precise and specific lineage tracing system for AdvSca1-SM cells^21^. Using the *Gli1* model, we found that, in response to acute vascular injury, AdvSca1-SM cells are rapidly activated, differentiate into myofibroblasts, and are major contributors to vascular remodeling and fibrosis; this cell transition is associated with loss of KLF4^20,21^. More recently, we reported that AdvSca1-SM cells modulate atherosclerotic plaque composition and *Klf4* depletion in AdvSca1-SM cells contributes to a more stable plaque phenotype^22^. Similarly, the *Gli1* lineage tracing model was used to show that a population of perivascular *Gli1*^+^ “mesenchymal stem cell (MSC)-like cells” are the major source of profibrotic cell populations, and that specific genetic ablation of these cells reduced organ fibrosis and injury^23,24^. However, detailed mechanisms regulating their stemlike phenotype and profibrotic transitions are unknown. In addition, the origin of *Gli1*-expressing cells in cardiac tissue and their relationship to AdvSca1-SM cells remain undefined.

In the current study, we used the *Gli1*-driven lineage tracing system to examine the phenotype and differentiation trajectory of coronary artery AdvSca1-SM cells in the setting of pressure overload-induced cardiac hypertrophy and fibrosis. Our data support that coronary SMC-derived AdvSca1-SM cells adopt a KLF4-dependent myofibroblast phenotype and are major contributors to cardiac fibrosis and inflammation. Importantly, the AdvSca1-SM-to-myofibroblast differentiation is observed in human cardiac hypertrophy, strongly supporting the translational significance of this cell population in human heart diseases.

## METHODS

### Mice

*Gli1*-Cre^ERT^ (JAX stock 007913) and ROSA26-YFP reporter mice (JAX stock 006148) were obtained from Jackson Laboratory. *Klf4* floxed mice were obtained from Dr. Klaus H. Kaestner (University of Pennsylvania). All mice were fully backcrossed to a C57BL/6 genetic background prior to studies. *Gli1*-Cre^ERT^ transgenic mice and Rosa26-YFP reporter mice were bred to generate tamoxifen-inducible AdvSca1-SM cell-specific YFP-expressing reporter mice (*Gli1*-Cre^ERT^-YFP). *Gli1*-Cre^ERT^-YFP mice were bred to *Klf4* floxed mice to generate AdvSca1-SM cell-specific *Klf4* knockout mice (*Gli1*-Cre^ERT^-YFP;*Klf4*^flox/flox^). 8-week old *Gli1*-Cre^ERT^-YFP mice received 1 mg IP tamoxifen injections for 12 consecutive days to induce YFP reporter knock-in and *Klf4* knockout prior to experiments. The tamoxifen was allowed to wash out for 5 days before the start of Angiotensin II or saline infusion. All surgeries were performed in a clean environment with sterile instruments. Micro-osmotic pumps (ALZET Model 1002 and Model 1004) were filled with calculated concentrations of Angiotensin II (AngII) or saline adjusted to the body weight of each individual mouse to ensure the delivery of AngII at the rate of 1 µg/kg/min per manufacturer instructions. The pumps were primed in sterile saline at 37°C for 24 hours. Primed pumps were inserted subcutaneously through a small incision at the dorsal neck region of the isoflurane-anesthetized mice and the incision was closed with stainless steel wound clips (CellPoint Scientific). After 14 days or 28 days of treatment, the mice were euthanized with isoflurane and cardiac tissues were dissected and processed for different assays accordingly. Both male and female mice were used for all studies. Mice were maintained in the Center for Comparative Medicine, and procedures were performed under a protocol approved by the Institutional Animal Care and Use Committee at the University of Colorado Denver.

### Histochemical fibrosis staining

Harvested aortic tissues were fixed in 10% buffered formalin phosphate (Fisher Scientific) for 48 hours and stored in 70% ethanol for 12 hours. Fixed tissues were processed with ASP300 S Enclosed Tissue Processor (Leica), embedded in paraffin and sectioned with a microtome at 5µm thickness. Masson’s trichrome staining was performed at the Histology Shared Resource core facility at the University of Colorado Cancer Center. Bright field images were obtained with an Olympus microscope at 20x magnification with SPOT software. All imaging was performed using the same setting.

### Immunofluorescent staining and analysis

Harvested cardiac tissues were fixed in 4% paraformaldehyde for 24 hours and cryopreserved in 30% sucrose for 24 hours. Tissues were embedded in OCT compound (Fisher HealthCare) and stored at −80°C. Tissues were sectioned at 6µm thickness with a cryostat. Sections were incubated with a rabbit monoclonal anti-GFP antibody (1:200, Abcam, Cat #Ab290) and a rat anti-CD68 antibody (1:100; Bio-Rad, Cat #MCA1957) overnight at 4°C followed by corresponding Alexa Fluor 488- and Alexa Fluor 594-conjugated secondary antibodies (1:500; Invitrogen) or anti-αSMA-Cy3 antibody (Sigma Aldrich, Cat #C6198, 1:2000) for 60 minutes at room temperature. Sections were mounted with Vectashield mounting media with DAPI (Vector Laboratories, Cat #H-1200). 40x or 60x images were captured with a Keyence BZ-X710 fluorescent microscope. Cellpose3 was used for fluorescent image segmentation. The pretrained model CP was further trained with a set of randomly selected 4 images. All images were segmented using the custom model and identified cells were exported as imageJ ROIs with the parameter --save_rois. ImageJ macro was used to quantify green and red fluorescence in all the ROIs. Custom python code was used to identify YFP^+^ and CD68^+^ cells. The identified positive cells were exported as ImageJ ROI and visually examined to rule out any false positive due to autofluorescence. Final quantification results were averaged per sample and expressed as cell count per 40x field.

### Second harmonic generation (SHG) imaging and analysis

Tissue sections were stained for YFP as described above to identify YFP^+^ AdvSca1-SM and AdvSca1-SM-derived cells and imaged for YFP and label-free SHG using a laser-scanning confocal microscope (LSM 780) at the UCAMC Advanced Light Microscopy Core facility, as described previously^21^. SHG signal on the red channel was quantified using ImageJ.

### Single cell RNA sequencing and data analysis

Cardiac tissues were harvested from *Gli1*-Cre^ERT^-YFP mice or *Gli1*-Cre^ERT^-YFP;*Klf4*^flox/flox^ treated with AngII or saline for 14 days. A Langendorff-Free approach^25^ was employed during the harvest to digest the cardiac tissue *in situ*. Briefly, mice were pretreated with an IP injection of 200 U heparin (Sigma, H3393), and euthanized with isoflurane 30 minutes later. Cardiac tissues were exposed, the inferior vena cava and descending aorta were cut, and 7 mL of EDTA buffer (113 mM NaCl, 5 mM KCl, 0.5 mM NaH_2_PO_4_ 10 mM HEPES, 5 mM EDTA, 10 mM glucose, 10 mM 2,3-Butanedione 2-monoxime (BDM), 10 mM Taurine, PH 7.8) was perfused from the right ventricle. The aortic arch was clamped and 10 mL of EDTA buffer was perfused from the left ventricle over 6 min, followed by 3 mL of perfusion buffer (113 mM NaCl, 5 mM KCl, 0.5 mM NaH_2_PO_4_ 10 mM HEPES, 1 mM MgCl_2_, 10 mM glucose, 10 mM 2,3-Butanedione 2-monoxime (BDM), 10 mM Taurine, PH 7.8) and finally 30-60 mL warm digestion buffer (3.2 mg/mL collagenase II, 0.7 mg/mL elastase, 0.2 mg/mL soybean trypsin inhibitor), until the cardiac tissues were fully digested. Cardiac tissues were further physically disrupted using a 10 mL serological pipet in 5mL of digestion buffer. Cell suspension was filtered through a 250 μm filter, washed once with FA3 buffer (1x PBS, 1mM EDTA, 25mM HEPES (pH 7.0), 1% FBS) and filtered through a 70 μm filter. Cell suspensions were pelleted and resuspended in 500 uL of FA3 buffer. The cell suspension was labelled with an anti-CD31-APC antibody (Invitrogen Cat #17-0311-80, 1:100) at 4 °C for 1 hour. Fluorescence activated cell sorting (FACS) was performed to separate CD31^+^ endothelial cell from the rest of the cells. For the scRNA-seq experiment comparing *Klf4* WT and KO mice, YFP^+^ AdvSca1-SM lineage cells were also separated. Sorted cells from two animals were pooled together for sequencing. The sorted CD31^+^ endothelial cells were mixed back with non-endothelial cells at 1:9 ratio, to avoid oversampling of the abundant endothelial cells^7^. For the experiment comparing YFP fluorescent positive and transcript positive cells, YFP^+^ cells were separated and sequenced separately. For scRNA-seq comparing *Klf4* WT and KO samples, YFP^+^, CD31^+^ and the YFP^−^ CD31^−^ populations mixed at 60:4:36 ratio, to increase the sampling of AdvSca1-SM cells in the final data. Samples were subjected to scRNA-seq using the Chromium Single Cell 3’ Library and Gel Bead Kit (v3.1 10x Genomics, Pleasanton, CA) and the Chromium X. Libraries were sequenced on an Illumina Novaseq 6000 at Genomics Shared Resource at the University of Colorado Anschutz Medical Campus. 50,000 reads per cell were obtained. Fastq files were aligned to custom build GRCm39 reference (Ensembl, r104) with the addition of YFP gene, using Cell Ranger 6.1.2. Scanpy 1.9.3 ^26^ was used for the downstream analysis including quality control, normalization, clustering, based on Single-cell best practices ^27^. Cells with less than 200 genes and genes expressed by less than 10 cells were filtered out. Cells with a percentage of mitochondrial counts exceeding 8 % or with more than 7500 genes were filtered. Pseudobulk profiles were created from the single-cell data using decoupler 1.6.0 ^28^. Differential gene expression analysis was performed with python implementation of the framework DESeq2^29^. Functional enrichment of biological terms was performed with decoupler 1.6.0 using GO_Biological_Process_2021.gmt, Reactome_2022.gmt, and KEGG_2019_Mouse.gmt obtained from Enrichr gene-set library^30^. Transcription factor activity inference was performed with decoupler 1.6.0 using CollecTRI gene regulatory network data^31^.

### Spatial transcriptomics

Spatial transcriptomics experiment was performed based on an official protocol from 10x Genomics. Briefly, cardiac tissues from 4-week AngII or saline treated mice were harvested and embedded in OCT in an isopentane and liquid nitrogen bath. 10 μm sections were prepared and placed onto the Visium Spatial Tissue Optimization Slide (PN-3000394) or Visium Spatial Gene Expression Slide (PN-2000233). Tissue permeabilization optimizations were performed and 12 min was determined to be suitable for mouse cardiac tissue. Tissue sections on Visium Spatial Gene Expression Slide were fixed with methanol at −20 °C for 30 min and labelled with an anti-αSMA antibody (Cy3 conjugated, 1:2000, Sigma C6198) for 30 min at room temperature followed by DAPI nuclear counter stain. 20x images of the whole tissue and fiducial frames were obtained through the XY stitching function of the Keyence BZ-X700 microscope. Once the image capture was completed, tissues were permeabilizated and reverse transcription, second strand synthesis and denaturation were performed. The samples were submitted to the Genomics Shared Resource core facility for cDNA amplifications and sequencing. Manual image alignment was performed with Loupe Browser. Space Ranger was used to process the fastq files with manually aligned immunofluorescent images of αSMA and DAPI. Downstream QC, analysis and visualizations were performed with Scanpy 1.9.3 and Squidpy 1.2.2^32^. Cell2location 0.1.3^33^ was used to deconvolute the spatial transcriptomics data by integrating with our scRNA-seq data. The composition information was used to cluster each tissue spots.

### Bulk RNA sequencing and data analysis

RNA-seq was performed as described previously^21^. In brief, single cell suspensions were prepared, as described above. FACS was performed to sort YFP^+^Sca1^+^ cells. Total RNA was extracted from FACS–sorted cells using QIAshredders and a RNeasy Plus Micro kit (QIAGEN). RNA-Seq library preparation and sequencing were conducted at the Genomics and Microarray Core at the University of Colorado Anschutz Medical Campus. Libraries were constructed using a SMARTer Stranded Total RNA-Seq kit (Clontech). Directional mRNA-Seq of sorted cell populations was conducted using the Illumina HiSeq 6000 system. HTStream 1.3.3 (https://github.com/s4hts/HTStream) was used to remove adaptor and rRNA sequence. Fastq files were aligned Mus musculus GRCm39 reference genome (Ensemble, release 104) using STAR 2.7.10b^34^ with default settings. Morpheus (https://software.broadinstitute.org/morpheus) was used to create heatmaps.

### Fixed RNA profiling

Cardiac tissues from patients with cardiomyopathy were dissected and segments of coronary arteries 5-7 cm in length were collected, minced, and digested into single cell suspension using digestion buffer (3.2 mg/mL collagenase II, 0.7 mg/mL elastase, 0.2 mg/mL soybean trypsin inhibitor), as previously published^21^. Cell suspensions were stained with DAPI viability dye and DAPI negative cells were collected using FACS (Sony MA900). The sorted viable cells were pelleted and resuspended in 1 mL fixation buffer (1x Fix & Perm Buffer(10x Genomics PN-2000517), 4% formaldehyde) and incubated at 4 °C for 20 hours. Fixed cells where centrifuged at 850 rcf for 5 min and pellets were resuspended in 1 mL of chilled quenching buffer (10x Genomics PN-2000516). 100 ul of pre-warmed enhancer (10x Genomics PN-2000482) and 275 uL 50% glycerol was added to the suspension and the samples were stored at −20 °C until submission to Genomics Shared Resource. Single cell suspensions were sequenced as described above.

### GTEx data analysis

Data were downloaded from the Genotype-Tissue Expression (GTEx) project and the samples were ranked by their Natriuretic Peptide B (NPPB) expression levels (NPPB is a cardiac fetal gene associated with pathological cardiac hypertrophy). 30% of subjects with highest and lowest NPPB expression in each sex and age groups were designated as hypertrophic and non-hypertrophic samples, respectively. Lima-Voom was used to examine the differentially expressed genes. RNA-seq data from corresponding arterial tissues were also examined.

### Data availability

The NCBI Gene Expression Omnibus database accession number for the data reported in this paper is GSE262589.

### *In vitro* AdvSca1-SM cell culture

*Gli1*-Cre^ERT^-YFP mouse and human coronary arteries were prepared into single cell suspensions, as described above. Mouse cells were labelled with an anti-Sca1-APC (eBioscience, Cat #17-5981-82) antibody, and human cells were labelled with anti-CD31-APC-Cy7 (BioLegend Cat #303119), anti-CD45-AF700 (BioLegend Cat #304023), and anti-CD34-PerCP-Cy5.5 (Biolegend Cat #343521)(the *Ly6a/*Sca-1 gene was deleted between mouse and rat speciation and, therefore, is not expressed in the human genome). FACS was performed using Sony MA900 sorter and YFP^+^Sca1^+^ cells were collected as mouse AdvSca1-SM cells while CD45^−^CD31^−^CD34^+^ cells were collected as AdvSca1-SM human equivalents. Sorted cells were plated in gelatin-coated plates with AdvSca1-SM media (α MEM [Gibco, Cat# 32571036]), 10% MSC qualified fetal bovine serum (FBS)(Thermofisher Cat #12662029), 1x Penicillin Streptomycin, 1ng/mL murine basic fibroblast growth factor (R&D systems 3139-FB), and 5ng/mL murine epidermal growth factor (R&D systems 2028-EG) at the density of 20,000/cm^2^. Cells used for experiments did not exceed passage 8.

### Quantitative RT-PCR

Total RNA was isolated from cultured cells by resuspending in RLT lysis buffer (Qiagen). Samples were then processed with QIAshredder and RNeasy Plus kits (Qiagen) to isolate total RNA. First strand cDNA was made using the qScript XLT cDNA SuperMix synthesis kit (Quantabio). Sequence-specific primers were designed (**Supplemental Table 1**). Quantitative real-time PCR (qPCR) was performed as previously described and 18S was used for normalization.

### Statistics

All experiments reported were carried out with at least 3 biological replicates, including both male and female mice. We found no differences between males and females; therefore, the data were combined for analysis. Data were analyzed using GraphPad Prism 10 (GraphPad Software, Inc). Shapiro-Wilk test (n≥3) or D’Agostino & Pearson test (n≥8) were performed to determine the normality of the data. Brown-Forsythe test was used to examine the equality of group variances. One-way ANOVA with Bonferroni’s post-hoc test was used to compare between multiple groups. For data that failed the normality test or equal variances test, a Kruskal-Wallis test was used to compare the groups followed by Dunn’s multiple comparison tests. Histochemical- and immunofluorescently-stained sections were visually examined and only sections with high quality tissue and staining were included in the quantitative analysis. P-values <0.05 were considered statistically significant.

## RESULTS

### Cardiac YFP^+^ cells from *Gli1*-CreERT-YFP mice exhibit high levels of smooth muscle cell-specific H3K4Me2 epigenetic lineage mark

Our previous data used chromatin immunoprecipitation (ChIP) approaches for a specific SMC epigenetic lineage mark as a secondary measure of the SMC origin of vascular AdvSca1-SM cells^20^. To define the SMC origin of *Gli1*^+^ lineage cells in cardiac tissue, we isolated AdvSca1-SM cells (YFP^+^Sca1^+^) cells, SMCs (YFP^−^Cd31^−^CD45^−^MCAM^+^), and non-SMCs (YFP^−^Cd31^−^CD45^−^ MCAM^−^)(**Supplemental Figure IA**) from single cell suspensions prepared from *Gli1-*Cre^ERT^-YFP mouse cardiac tissue using Fluorescence-activated Cell Sorting (FACS). The sorted cell populations were subjected to ChIP analysis using an anti-H3K4Me2 antibody, a well-recognized chromatin modification observed on histones flanking CArG elements of contractile gene promoters and highly specific to SMC lineage^20,35^. Similar to cardiac SMCs, cardiac AdvSca1-SM cells exhibited high levels of the H3K4Me2 mark on histones flanking CArG elements of both the SMC-specific *Acta2* and *Myh11* gene promoters (**Supplemental Figure I B&C**, respectively), strongly supporting the SMC origin of *Gli1* expressing cells in cardiac tissue. Combined with our previous publications^20,21^, this data confirms that *Gli1*-driven reporter mice are a faithful lineage tracing system of AdvSca1-SM cells in the cardiovascular system.

### Perivascular coronary artery AdvSca1-SM cells expand and infiltrate into the cardiac interstitium surrounded by and embedded in a rich collagen matrix

To examine the response of AdvSca1-SM cells in pathological cardiac fibrosis, *Gli1-*Cre^ERT^-YFP mice were subjected to Angiotensin II (AngII)-induced cardiac hypertrophy and fibrosis^36^. Compared to saline controls, 4-week AngII treatment promoted extensive perivascular and interstitial fibrosis (Masson’s trichome staining and second harmonic generation [SHG] imaging, **Figure 1A&B, Supplemental Figure II)**. To examine the spatial relationship between AdvSca1-SM cells and cardiac fibrosis, we further visualized YFP^+^ AdvSca1-SM cells overlaying the SHG images (**Figure 1B; Supplemental Figure II**). αSMA staining was used to identify normal (saline) and hypertrophic (AngII) coronary arteries (**Figure 1C**). Results demonstrated that, similar to our previous data in the vasculature, in control mice (saline), AdvSca1-SM cells were localized to a perivascular adventitial niche underlying the vascular media. In contrast, in response to AngII, AdvSca1-SM cells expanded in number within the adventitia and migrated from a perivascular coronary artery adventitial niche into the cardiac interstitium embedded in and surrounded by a rich fibrillar collagen matrix, consistent with AngII-mediated cardiac fibrosis (**Figure 1B&C; Supplemental Figure II**).

**Figure 1.**
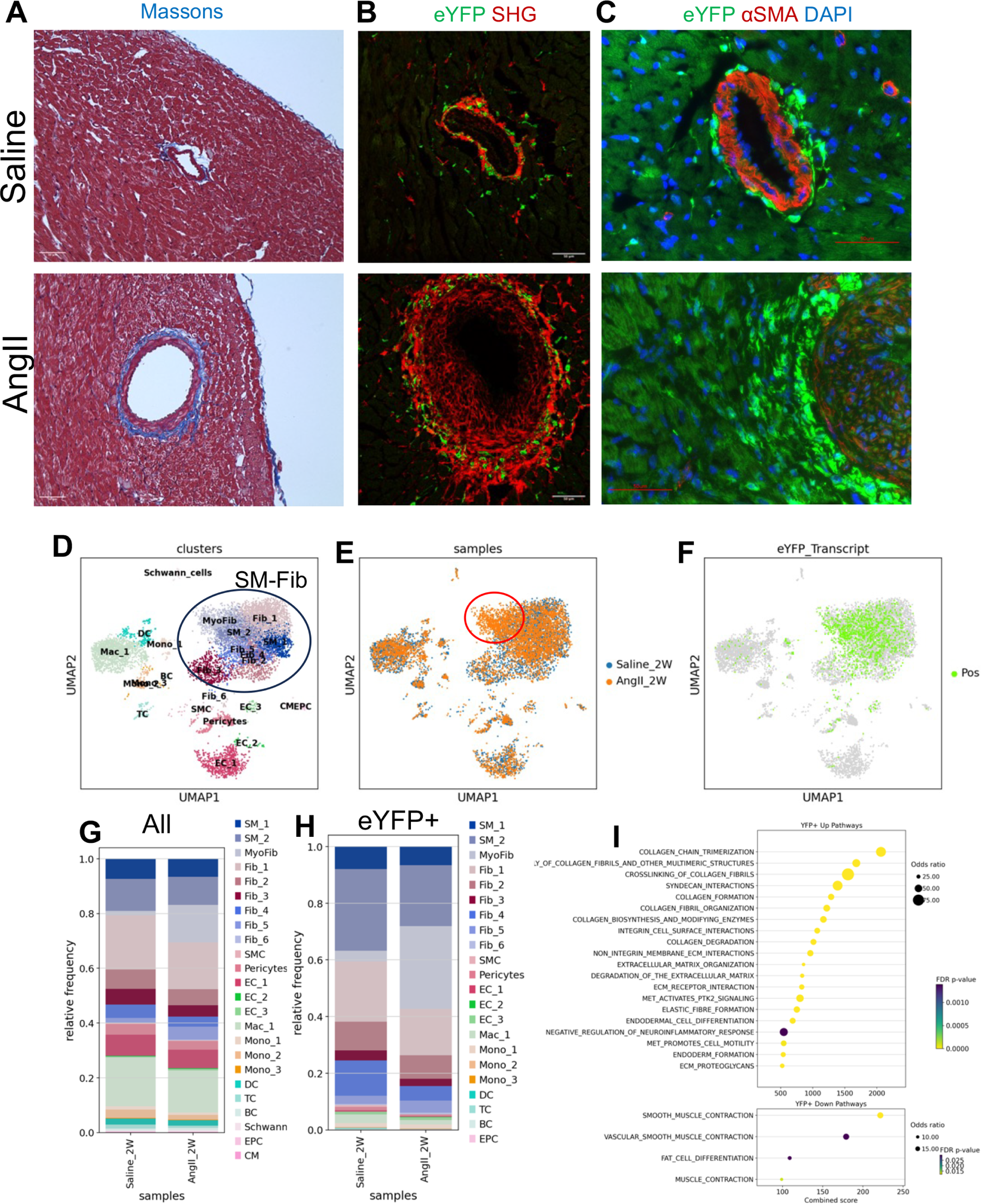
Perivascular AdvSca1-SM cells expand, infiltrate, and adopt a myofibroblast phenotype to contribute to pathological cardiac fibrosis. *Gli1*-Cre^ERT^-YFP mice were subjected to Angiotensin II (AngII)-induced cardiac hypertrophy and fibrosis, as described in the Methods section. **(A)** Masson’s trichrome staining of 4-week AngII or saline treated cardiac tissue; ECM/collagen deposition shown in blue. Representative 20x images of N=4 in each group are shown. **(B)** Cardiac tissue sections were immunofluorescently stained with anti-GFP antibody and subjected to label-free SHG imaging to visualize the collagen deposition (Red). YFP^+^ cells were imaged (Green) and overlayed to examine their association with ECM/collagen deposition. Representative 40x images of saline (N=10) and AngII-treated (N=12) tissues are shown. (**C**) Cardiac tissue sections were immunofluorescently stained with anti-GFP (Green), anti-αSMA (Red) and DAPI. Representative 60x images of N=6 in each group are shown. **(D-I)** Cardiac tissue from 2-week AngII- or saline-treated mice were harvested and prepared for scRNA-seq, as described in the Methods section. **(D)** Uniform Manifold Approximation and Projection (UMAP) visualization is shown. Blue circle highlights the major cluster that is annotated as AdvSca1-SM and fibroblast (SM-Fib) clusters. **(E)** scRNA-seq data UMAP plot colored by treatment (Saline - blue; AngII - orange). Red circle highlights the AngII-induced shift. **(F)** YFP transcript positive cells were highlighted in green in the UMAP plot. **(G&H)** Composition of Saline and AngII-treated samples were visualized for all cells **(G)** and YFP^+^ cells **(H)** of the scRNA-seq data. **(I)** Pathway analysis for genes up-regulated (top) and down-regulated (bottom) by AngII treatment in YFP^+^ AdvSca1-SM cells.

### AdvSca1-SM cells adopt a myofibroblast phenotype in response to AngII-induced cardiac fibrosis

To better understand the transcriptomic changes of cardiac cells, including AdvSca1-SM cells, in the setting of cardiac fibrosis, we performed scRNA-Seq of single cell suspensions prepared from heart tissues from saline- vs AngII-treated mice, as described in the method section (**Supplemental Figure III**); cardiomyocytes were excluded due to cell size. Heart cell types were annotated by their gene expression profiles (**Figure 1D, Supplemental Figure IVA**). A large cluster consisted of multiple subclusters that expressed AdvSca1-SM (SM_1 and SM_2), fibroblast (Fib1-5), and myofibroblast (MyoFib) gene signatures (**Figure 1D; SM-Fib cluster**). Compared to saline controls, AngII-treated hearts exhibited an expansion of myofibroblasts within the SM-Fib cluster (**Figure 1E, red circle &F**). Overall, Differential gene expression (DGE) analysis (**Supplemental Table 2**) and pathway analysis of the SM-Fib cluster revealed that AngII stimulated a pro-fibrotic response of all SM-Fib cells (**Supplemental Figure IVB, Supplemental Table 3**). Macrophages and monocytes (DGE analysis)(**Supplemental Table 4**) showed elevated signatures of recruitment and proinflammatory activation in response to AngII (**Supplemental Figure IVC, Supplemental Table 5**). We selectively examined the AdvSca1-SM cells and AdvSca1-SM-derived cells in the scRNA-seq data identified by YFP mRNA expression. YFP-expressing AdvSca1-SM cells predominantly clustered within the major SM-Fib cluster, supporting their contribution to ECM deposition, with rare contributions to SMCs, pericytes, endothelial cells, and monocytes and macrophages (**Figure 1F**), consistent with our previous data demonstrating the multipotent ability of AdvSca1-SM stem cells to differentiate into multiple cell types^20^. Composition analysis showed that 29.2% of **SM-Fib** cells adopt a myofibroblast phenotype in response to AngII treatment, a sharp contrast to 3.7% in saline treated hearts (**Figure 1G&H**). Interestingly, all YFP^+^ cells within the major SM-Fib cluster, including AdvSca1-SM cells in both saline and AngII conditions, expressed the transcription factor, *Tcf21* (**Supplemental Figure V**), consistent with a previous manuscript describing resident cardiac fibroblasts from the *Tcf21* lineage^37^. DGE analysis (**Supplemental Table 6**) and pathway analysis showed that AngII stimulation induced a profibrotic transition of YFP^+^ cells (**Figure 1I top, Supplemental Table 7**) and suppressed genes related to smooth muscle contraction (**Figure 1I bottom, Supplemental Table 8**).

We performed a trajectory analysis of the SM-Fib cells in the scRNA-seq data. RNA velocity analysis revealed a differentiation trajectory along SM_1, SM_2, and MyoFib clusters (SM-to-MyoFib trajectory)(**Figure 2A**). Pseudotime analysis confirmed that SM_1 and SM_2 clusters represent earlier timepoints in the differentiation trajectory to MyoFib cluster (**Figure 2B**). We further used CellRank to estimate cell fate probability among the SM-Fib cells. While most of the cells were predicted to be on the trajectory towards the MyoFib cluster (**Figure 2C**), the algorithm identified a separate trajectory mostly contained within Fib3 (**Figure 2D**). Cell cycle analysis showed a gradual increase in cell proliferation along the SM-to-MyoFib differentiation trajectory, which was not observed in the Fib3 trajectory (**Figure 2E**). We examined the trend of select gene expression along the SM-to-MyoFib trajectory. Associated the with trajectory, there is a gradual decrease of expression of genes related to AdvSca1-SM cell stemness (*Ly6a*, *Ly6c1*, *Cd34*, *Scara5*, *Ackr3*, *Klf4*; **Figure 2F**) and an increase of profibrotic genes (*Postn*, *Fn1*, *Col1a1*, *Col1a2*, *Col3a1*, *Runx1*; **Figure 2G**), supporting that AdvSca1-SM cells lose their progenitor phenotype and differentiate towards myofibroblasts. DGE (**Supplemental Table 9**) and pathway analysis (**Supplemental Table 10**) comparing YFP^+^ MyoFib clusters and SM clusters confirmed the up-regulation of ECM related pathways (**Supplemental Figure IVD**) and down-regulation of stemness related pathways such as vasculogenesis (**Supplemental Figure IVE, Supplemental Table 11**). Transcription factor (TF) activity was inferred based on transcriptomic data. The results showed MyoFib cluster exhibited an up-regulation of fibrosis-related TFs (TCF21, SMAD3, SMAD4) and down-regulation of a transcription factor that suppresses fibrosis (SMAD7)(**Supplemental Figure IVF**).

**Figure 2.**
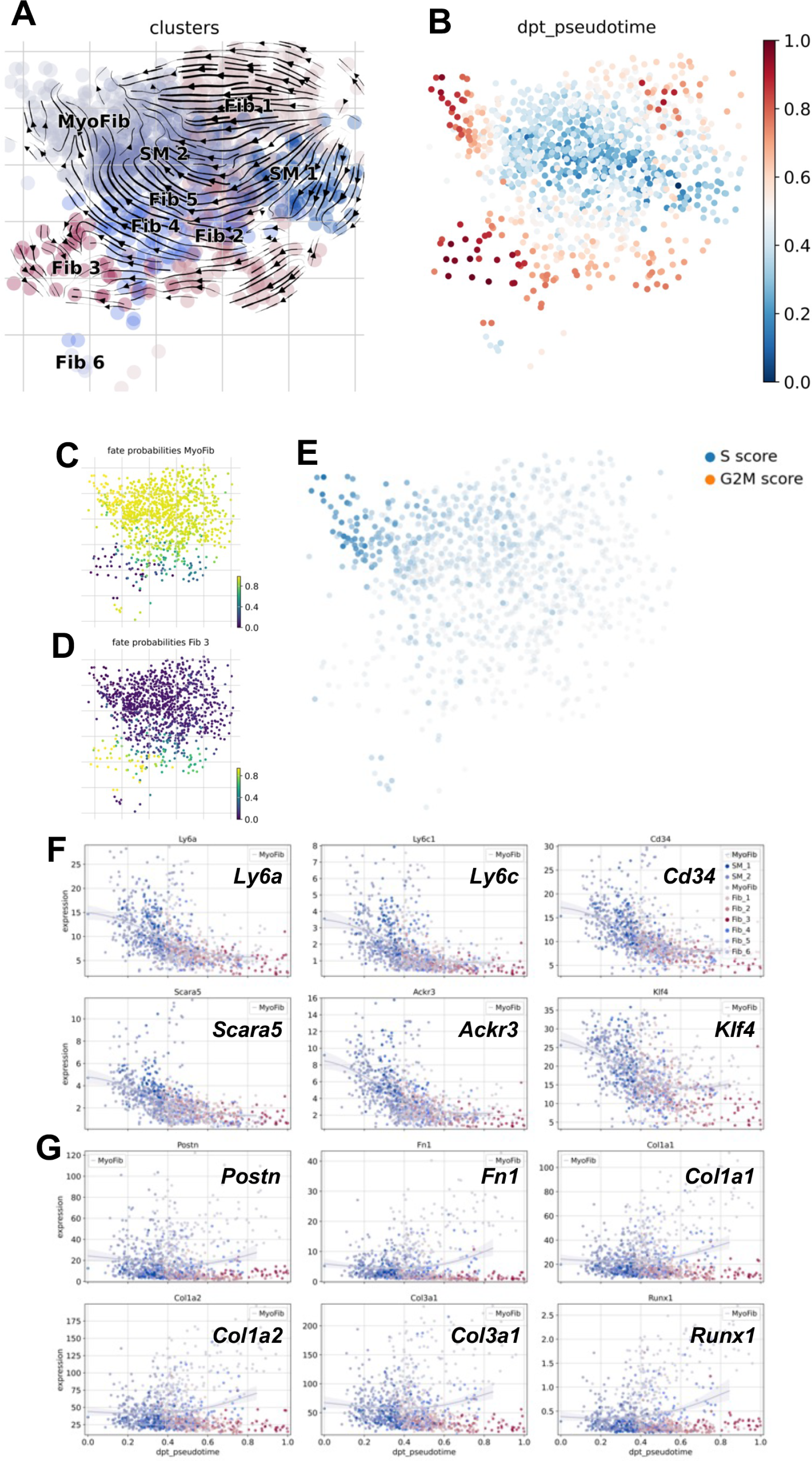
AdvSca1-SM cells adopt a profibrotic differentiation trajectory in response to AngII. **(A)** RNA velocity analysis was performed to visualize the differentiation trajectory of the SM-Fib subclusters of scRNA-seq data. Stream plot of RNA velocity embedded in the UMAP is shown. The length of each vector represents RNA velocity along the trajectory, and the width is proportional to the number of cells at a given position. **(B)** UMAP of SM-Fib subclusters is shown colored by diffusion pseudotime (dpt), where blue represents early in the trajectory and red represents late pseudotime. **(C&D)** CellRank was used to examine the fate probabilities of SM-Fib cells. The UMAP was colored by the probability of adopting a MyoFib fate **(C)** and a Fib3 fate **(D)**. **(E)** SM-Fib cells were scored for marker genes of S and G2M phase. UMAP is shown with colored by respective score. **(F&G)** Expression trend of select stemness genes **(F)** and profibrotic genes **(G)** are visualized along the pseudotime trajectory.

As we relied on YFP transcript to identify AdvSca1-SM cells in the scRNA-seq analysis, we performed a separate experiment where we separated YFP^+^ AdvSca1-SM cells and YFP^−^ cells using FACS and sequenced the two populations separately. YFP^+^ cells identified by transcript exhibited a similar distribution and composition (**Supplemental Figure VI A&B)** to that of the YFP^+^ cells by fluorescence (**Supplemental Figure VI C&D**), supporting that the YFP transcript can be used to accurately identify YFP^+^ AdvSca1-SM cells in the analysis of the scRNA-seq data.

### Gene signature of AdvSca1-SM-to-MyoFib transition is observed in human cardiomyopathy

To investigate whether the observed myofibroblast differentiation of AdvSca1-SM cells is present in human cardiac fibrosis, we obtained RNA-Seq data from human left ventricular tissue from the GTEx project. Samples were ranked by their natriuretic peptide B (*NPPB*) expression level as *NPPB* is a cardiomyocyte gene associated with the fetal gene program and pathological cardiac hypertrophy^7,38^. 30% of subjects with the highest and the lowest *NPPB* expression were designated as hypertrophic and non-hypertrophic samples, respectively^7^ (**Figure 3A**). DGE analysis was performed (**Supplemental Table 12**. Hypertension induced significant down-regulation of AdvSca1-SM stemness-related genes (**Figure 3B&D**) and up-regulation of myofibroblast genes (**Figure 3C&D**), consistent with the gene signature observed in the SM-to-MyoFib differentiation trajectory induced by AngII. A similar trend was observed in arterial tissues of matching subjects (**Figure 3E**). In addition, we prepared single cell suspensions from coronary arteries of human cardiomyopathy patients for single cell Fixed RNA profiling (10x Genomics) to further examine the gene signature of human AdvSca1-SM-like cells. Data were annotated through reference mapping to our mouse scRNA-seq data based on orthologous genes (**Figure 3F**). Human coronary AdvSca1-SM-like cells expressed stemness genes including *CD34, SCARA5, ACKR3, PI16* and *KLF4* at high levels, while the MyoFib cluster selectively expressed ECM genes (**Figure 3G**) suggestive of the translational relevance of a profibrotic transition of AdvSca1-SM cells in cardiac cardiomyopathy and fibrosis.

**Figure 3.**
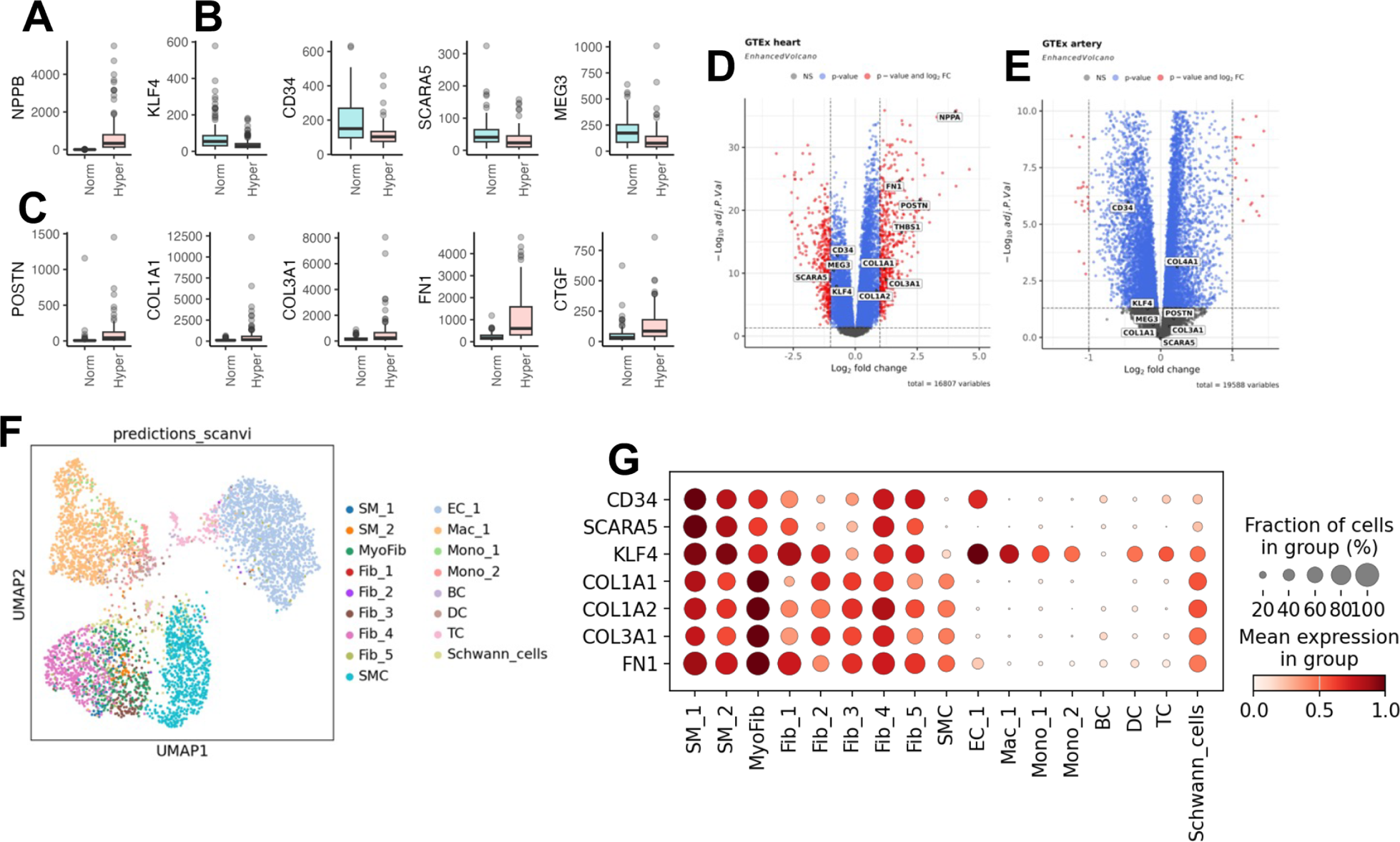
AdvSca1-SM-to-myofibroblast differentiation is observed in human hypertrophic cardiomyopathy cardiac tissue. RNA-seq data of left ventricular tissue were downloaded from the Genotype-Tissue Expression (GTEx) project and stratified and analyzed as described in the Methods section. **(A-C)** Differential gene expression (DGE) analysis was performed and normalized expression of *NPPB* **(A)** and select genes related to stemness **(B)** and fibrosis are shown **(C)**. **(D)** Volcano plot showing DGE of select genes related to AdvSca1-SM cell stemness and profibrotic transition are shown. Positive Log2FC indicate up-regulation in hypertrophic heart. **(E)** RNA-seq data of arterial tissue from corresponding subjects were retrieved; volcano plot is shown with select genes annotated. **(F)** Single cell suspensions were prepared from coronary arterial tissues collected from hypertrophic cardiomyopathy patients. Fixed RNA Profiling (10x Genomics) was performed to examine single-cell gene expression. UMAP plot is shown with annotation corresponding to clusters in mouse scRNA-seq data. **(G)** Select stemness and profibrotic genes expression are shown in dotplot.

### Spatial transcriptomics analysis revealed the spatial embedding of AdvSca1-SM cells and their relationship to cardiac fibrosis and other cell populations

To examine the spatial distribution of AdvSca1-SM cells in cardiac tissue, we used a spatial transcriptomics approach (Visium Spatial Gene Expression, 10x Genomics). Cardiac sections from AngII- or saline-treated mice were harvested and processed for spatial transcriptomics analysis, as described in the Methods Section. Cell2location was used to deconvolute the spatial transcriptomics data by integrating with the annotated scRNA-seq data (Figure 1C). Thirteen clusters were identified (**Figure 4A**) with AngII dramatically altering the clustering of the tissue spots as shown in the UMAP (**Figure 4B**) and the spatial embedding (**Figure 4C**). Strong YFP signature was observed in clusters 10 and 12 (**Figure 4D**), suggesting these clusters represent AdvSca1-SM and AdvSca1-SM-derived cells. Interestingly, cluster 12 exhibit high expression of AdvSca1-SM stemness genes while cluster 10 exhibited elevated myofibroblast genes expression (**Figure 4E**). In the visualization of the spatial tissue deconvolution, Clusters 10 and 12 exhibited high composition of cells corresponding to MyoFib and SM_1, respectively (**Figure 4F**), confirming their link to corresponding scRNA-seq clusters (Figure 1C). Limited by the resolution of the Visium platform, we could not establish that all cells covered by cluster 10 are myofibroblasts. Indeed, cluster 10 encompassed additional types of fibroblasts (**Figure 4F**). Interestingly, cluster 10 also exhibited an elevated composition of monocytes and T cells (**Figure 4F**), suggesting that YFP^+^ AdvSca1-SM-derived myofibroblasts are in close proximity to recruited immune cells and may play a vital role in the recruitment of immune cells and tissue inflammation in the setting of pathological cardiac fibrosis. Cluster 10 also exhibited the highest αSMA IF signal (**Figure 4G**), further supporting the conclusion that they represent myofibroblasts. AngII reduced Cluster 12 (SM_1, **Figure 4H**) and stimulated the expansion of cluster 10 (MyoFib) in perivascular and interstitial regions (**Figure 4I**). Together, the data mapped the spatial transition of AdvSca1-SM cells in response to AngII and confirmed their specific involvement in regions of fibrosis and inflammation.

**Figure 4.**
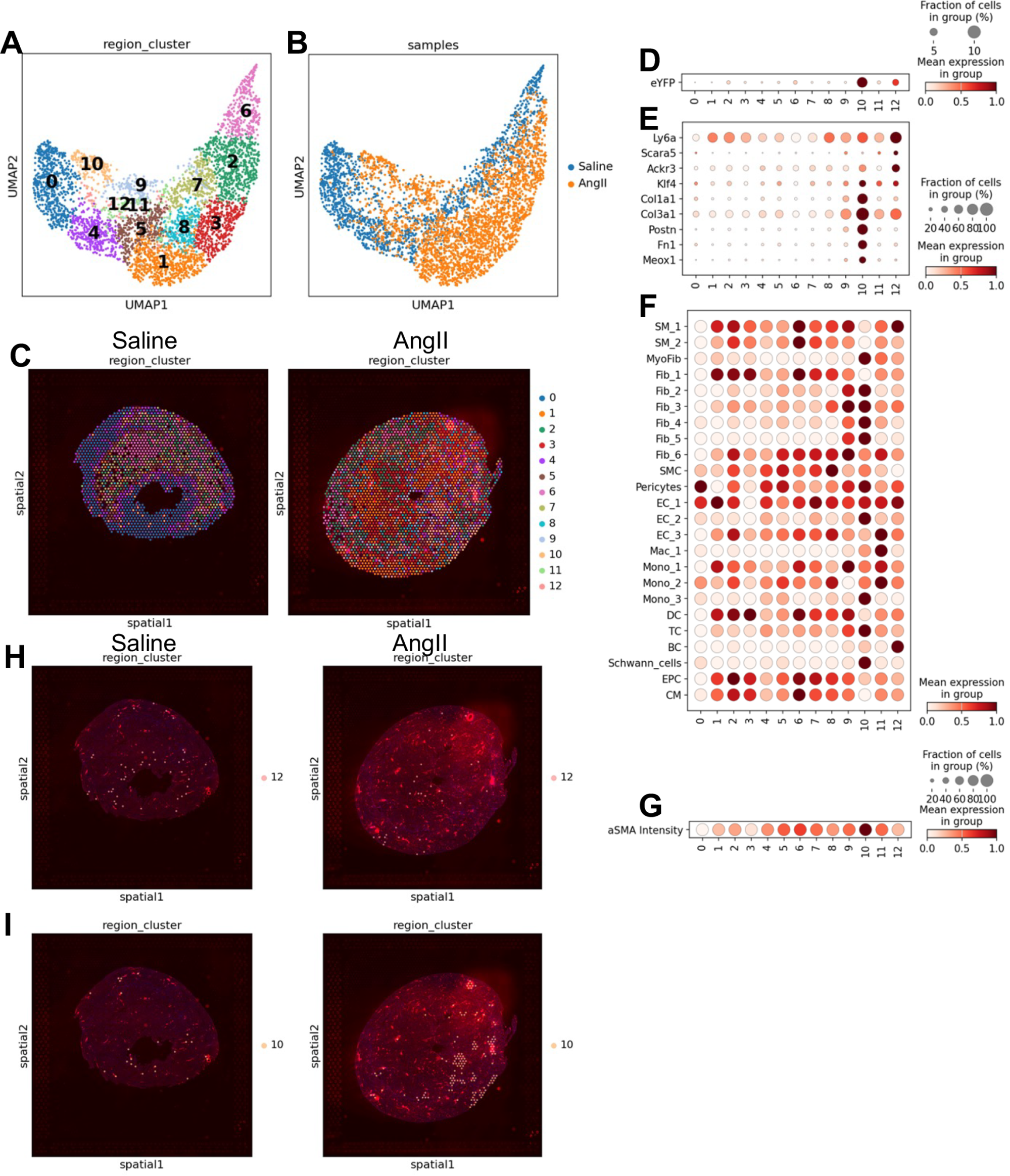
AngII-induced expansion and differentiation of AdvSca1-SM cells is revealed by spatial transcriptomics. Spatial transcriptomics was performed with cardiac tissue from AngII- or saline-treated cardiac tissue sections, as described in the Methods section. The data was deconvoluted using scRNA-seq data as reference. **(A)** UMAP plot showing the captured tissue spots clustered based on the deconvoluted data. **(B)** UMAP plot colored by saline- (blue) or AngII (orange)-treated tissues. **(C)** spatial embedding of the region clusters is shown for saline-treated (left) and AngII-treated (right) hearts. **(D&E)** Expression of YFP **(D)** and select stemness and profibrotic genes **(E)**, grouped by clusters, are shown in dotplot. **(F)** Dotplot showing the correlation between region clusters and scRNA-seq clusters are shown. Red color gradient corresponds to the composition of respective cell types in the region clusters. **(G)** α-SMA immunofluorescent signal intensity was quantified and shown in dotplot grouped by region clusters. **(H&I)** Spatial embedding of region cluster 12 **(H)** and cluster 10 **(I)** is highlighted and overlayed on the α-SMA IF staining of the tissues.

### KLF4 is an essential mediator of AdvSca1-SM cell proinflammatory and profibrotic response

We previously reported that loss of KLF4 expression is associated with decreased stemness in vascular AdvSca1-SM cells in the setting of vascular injury^21^. Our scRNA-seq data support that *Klf4* is highly expressed by cardiac AdvSca1-SM cells in their stemness state and repressed during the myofibroblast transition (Figure 2F & Figure 3); similar findings were observed in human cardiomyopathy compared to non-hypertrophic patients (Figure 3). To further study the role of KLF4 specifically in AdvSca1-SM cells, we established AdvSca1-SM-specific *Klf4* knockout (*Gli1*-Cre^ERT^-YFP;*Klf4*^flox/flox^ [*Klf4* KO]) mice by crossing AdvSca1-SM lineage mice with *Klf4*-floxed mice; tamoxifen treatment deletes KLF4 and activates YFP expression only in AdvSca1-SM cells. WT and *Klf4* KO mice were subjected to 4-weeks of AngII or saline treatment. Masson’s trichrome staining showed that AngII-induced perivascular cardiac fibrosis was blunted in *Klf4* KO mice (**Figure 5A**). SHG imaging was performed to quantify fibrillar collagen deposition. The results support that AngII-induced perivascular cardiac fibrosis was inhibited in *Klf4* KO mice (**Figure 5B**; quantified in **5C**). AngII-induced interstitial fibrosis was also blunted in *Klf4* KO mice, as evaluated by Masson’s trichrome staining (**Supplemental Figure VIIA**) and SHG (**Supplemental Figure VIIB**). Blunted fibrosis in *Klf4* KO mice was associated with reduced AngII-induced expansion of YFP^+^ AdvSca1-SM cells in *Klf4* KO mice (**Figure 5B**), indicating that *Klf4* is essential for the profibrotic response of AdvSca1-SM cells. AngII-induced cardiac hypertrophy, however, was not altered by AdvSca1-SM cell *Klf4* depletion (**Figure 5D**) demonstrating that the effects of AdvSca1-SM KLF4 are specific to pathological fibrosis and not pathological cardiac hypertrophy.

**Figure 5.**
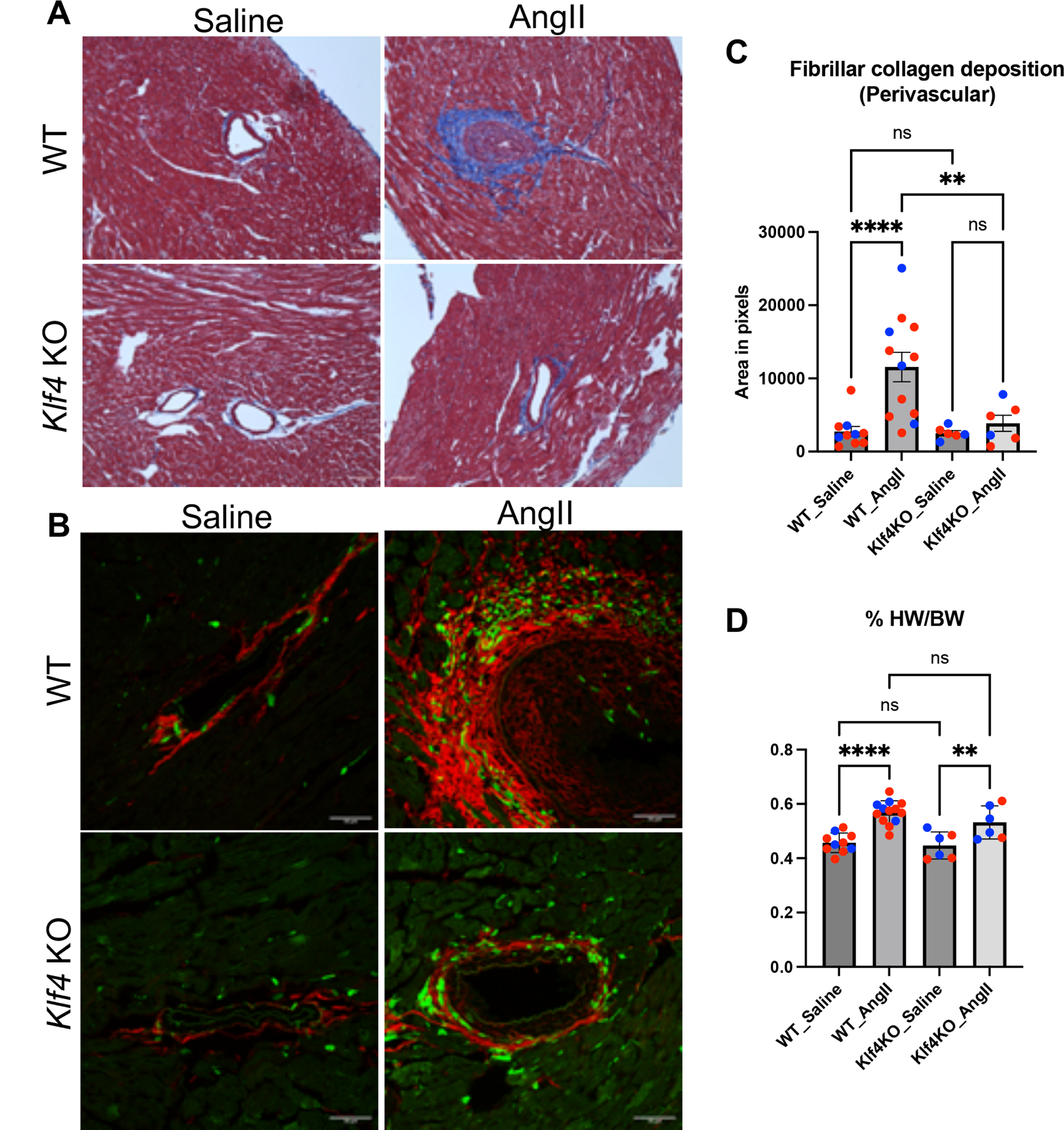
AngII-induced cardiac fibrosis is blunted in AdvSca1-SM-specific *Klf4* KO mice. WT and AdvSca1-SM-specific *Klf4* KO mice were subjected saline/AngII treatment for 4 weeks and the cardiac tissues were harvested. **(A)** Masson’s trichrome staining showing ECM deposition in perivascular regions. Representative 20x images of N=4 in each groups are shown. **(B)** SHG imaging (red) with YFP co-staining (green) was performed to show the spatial relation between AdvSca1-SM cells and ECM deposition in the differently treated mice. **(C)** Perivascular ECM content was quantified from the SHG images. **(D)** Normalized heart weight was shown in the different groups. Dots in panel C&D represent individual mice (blue: male; red: female).

To better understand the molecular function and signaling altered by AdvSvca1-SM *Klf4* depletion, scRNA-seq was performed with cardiac tissue from saline or AngII-treated WT and *Klf4* KO mice (**Figure 6A)**. Compared to WT mice, *Klf4* depletion did not introduce major changes in the overall composition of YFP^+^ and YFP^+^ SM-Fib cells in response to AngII (**Figure 6B**). DGE analysis comparing AngII-treated *Klf4* KO and WT YFP^+^ AdvSca1-SM cells surprisingly showed that *Klf4* KO AdvSca1-SM cells exhibited elevated expression of select collagens and matrix metallopeptidases (**Figure 6C, Supplemental Table 13**). Pathway analysis also supported that up-regulated genes in *Klf4* KO AdvSca1-SM cells are overrepresented in ECM-related pathways (**Figure 6D, Supplemental Table 14**). Notably, most of the up-regulated collagens are non-fibrillar collagens with the exception of *Col2a1*. In addition, some of the up-regulated pathways, such as “cardiac muscle tissue morphogenesis” and “regulation of cell growth involved in cardiac muscle cell development”, suggest the possibility that *Klf4* depletion promotes a AdvSca1-SM cell phenotype associated with cardiac tissue growth and repair. Compared to WT, *Klf4* KO AdvSca1-SM cells exhibited reduced levels of an array of cytokines and chemokines (**Figure 6C, Supplemental Table 13**). Accordingly, genes related to inflammation, and specifically interferon signaling, were suppressed in *Klf4* KO AdvSca1-SM cells (**Figure 6E, Supplemental Table 15**), indicating the protection against AngII-induced fibrosis observed in *Klf4* AdvSca1-SM KO mice is attributed to reduced pathological cardiac fibrosis and the anti-inflammatory phenotype of AdvSca1-SM cells. Further, analysis was performed to infer the TF activity based on the transcriptomics data altered by *Klf4* depletion in the setting of AngII treatment. TFs that promote inflammation and interferon signaling, including IRF1, IRF2, IRF7, NF-κB, and HIF1α are predicted to have lower activity in *Klf4* KO compared to WT AdvSca-1-SM cells (**Figure 6F**). Among the immune cell clusters in the scRNA-seq data, macrophages and monocytes (Mac-Mono) are the most prominent populations. We performed DGE analysis between Mac-Mono populations from AngII-treated WT and AdvSca1-SM *Klf4* KO hearts (**Figure 7A, Supplemental Table 16**). Mac-Mono populations from *Klf4* KO tissue exhibited reduced expression of interferon-related genes and pathways (**Figure 7A&B, Supplemental Table 17**) but *Klf4* deletion promoted a collagen and contractile phenotype of Mac-Mono clusters (**Figure 7C, Supplemental Table 18**). These data importantly suggest that manipulating the phenotype of AdvSca1-SM cells through *Klf4* depletion promotes a shift of macrophage phenotype from an inflammatory M1-like toward a reparative M2-like phenotype. To further examine the infiltration of macrophages in the cardiac tissue of AngII/saline treated WT and *Klf4* KO mice, cardiac tissue sections were stained for CD68^+^ macrophages. AngII-induced infiltration of macrophages in the perivascular regions was blunted in *Klf4* KO mice compared to WT mice (**Figure 7D**, quantified in **Figure 7E**), in association with the reduced expansion of YFP^+^ AdvSca1-SM cells (**Figure 7 D&E**). The suppression of inflammation and interferon signaling was also observed in endothelial cells (**Supplemental Figure VIIIA, Supplemental Table 19**) and smooth muscle cells/pericytes (**Supplemental Figure VIIIB, Supplemental Table 20**) in *Klf4* KO compared to WT mice, indicating AdvSca1-SM-specific *Klf4* depletion induces a systemic anti-inflammatory phenotype in resident cardiac and recruited inflammatory cells. To further validate these finding, we performed bulk RNA-seq with YFP^+^ AdvSca1-SM cells sorted from AngII or saline-treated WT and *Klf4* KO mice. Visualization of genes related to interferon pathways (Figure 6E) showed extensive induction in AngII-treated WT, but suppression in *Klf4* KO AdvSca1-SM cells (**Supplemental Figure VIIIC**).

**Figure 6.**
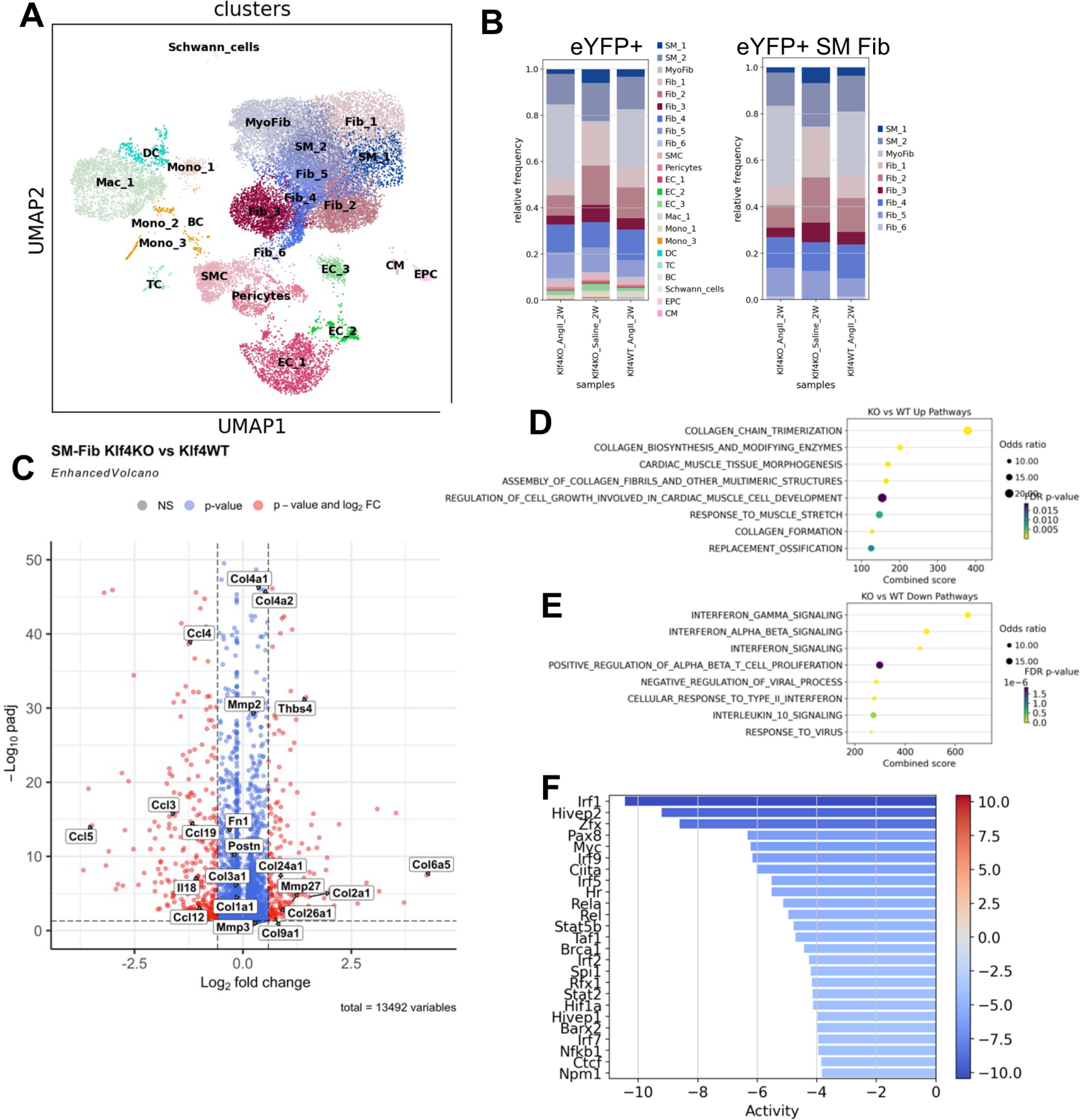
*Klf4* depletion in AdvSca1-SM cells elicits an anti-inflammatory phenotype. scRNA-seq was performed to examine gene expression profile of AngII/saline treated WT and *Klf4* KO cardiac tissue. UMAP **(A)** and composition analysis **(B)** of YFP^+^ cells are shown. **(C)** DGE analysis was performed to compare gene expression in YFP^+^ SM-Fib cells in AngII treated WT and *Klf4* KO heart. Select genes are shown in volcano plot. Pathway analysis was performed with up-regulated genes **(D)** and down-regulated genes **(E)** in *Klf4* KO YFP^+^ SM-Fib cells compared to WT. **(F)** Transcription factor activity was inferred based on the transcriptomic changes and the relative activity is shown.

**Figure 7.**
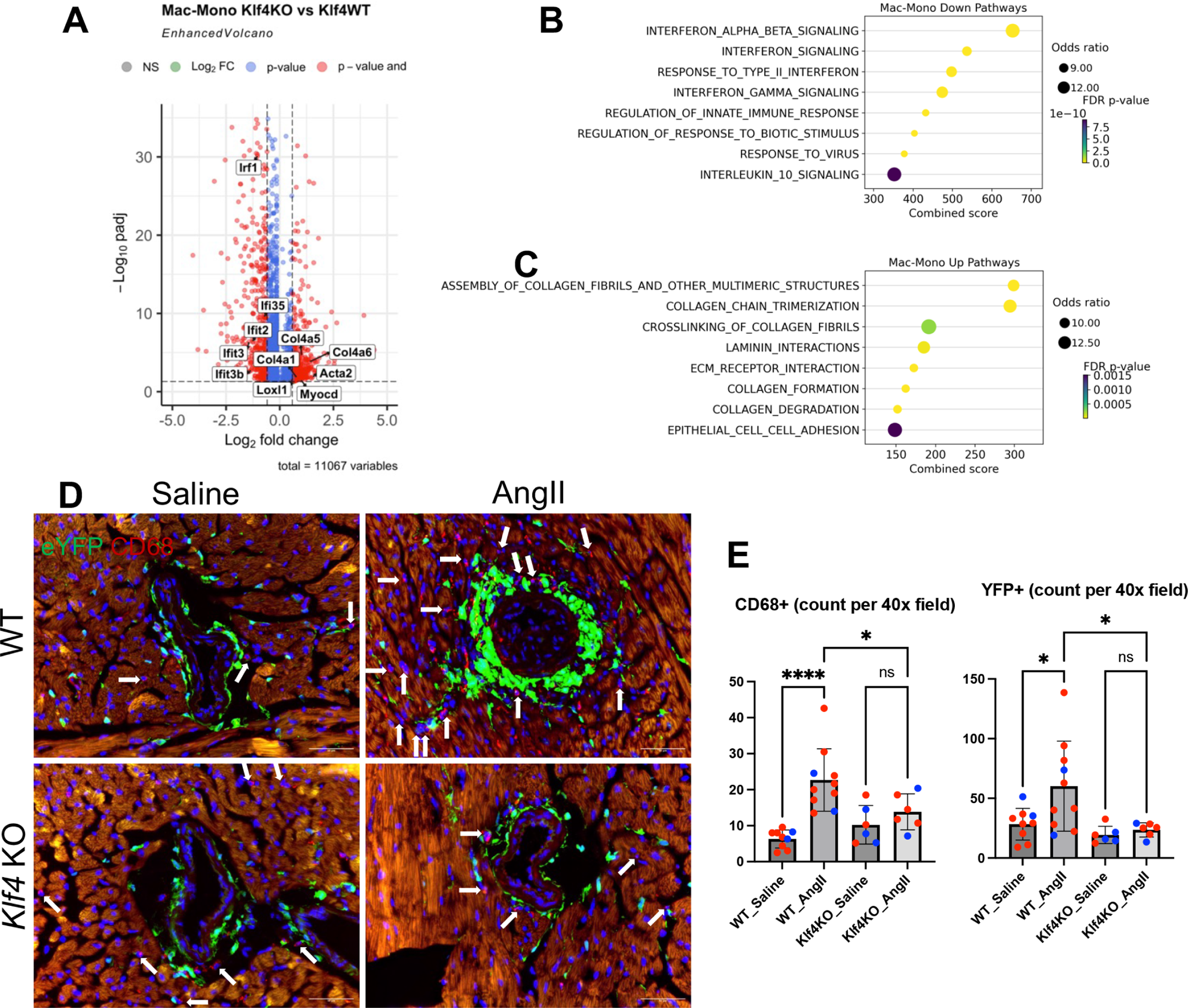
AdvSca1-SM cell *Klf4* depletion induce an anti-inflammatory phenotype in macrophages and defects in AngII-induced macrophage recruitment. DGE analysis was performed to compare macrophage and monocyte (Mac-Mono) gene expression profile in the scRNA-seq data. (A) select up-regulated and down-regulated genes in Mac-Mono cells in *Klf4* KO compared to WT heart are shown in volcano plot. Pathway analysis of down-regulated **(B)** and up-regulated **(C)** genes are shown. **(D)** Immunofluorescent staining for CD68^+^ macrophages was performed and representative images from AngII/Saline treated WT/*Klf4* KO heart are shown. Quantifications of CD68^+^ **(E)** and YFP^+^ AdvSca1-SM lineage cells **(F)** are shown. Dots in panel E&F represent individual mice (blue: male; red: female)

### KLF4 controls stemness and pro-fibrotic gene expression

To further examine the function of KLF4 in AdvSca1-SM cells, we established *in vitro* cultures of cardiac AdvSca1-SM (CSM) cells. CSM cells were treated with TGF-β for 24 hours as a pro-fibrotic stimulus^39^. Consistent with our previous report, stemness-related genes (*Klf4*, *Ly6a*, and *Cd34*) were suppressed by TGF-β treatment (**Figure 8A**). In addition, *Ackr3*, which encodes CXCR7, a chemokine receptor mediating vascular progenitor migration^40^, as well as the stem cell-associated lncRNA, *Meg3*^41^, were suppressed by TGFβ stimulation (**Figure 8A**). In contrast, expression of myofibroblast-related genes, including *Postn*, *Col1a2,* and *Col4a1* were induced by TGF-β treatment (**Figure 8A**). *In vitro* cultures of human AdvSca1-SM-like cells derived from donor coronary artery tissues (CD45^−^CD31^−^CD34^+^) were also established (**Supplemental Figure IXA**). Human AdvSca1-SM-like cells exhibited a similar response to TGF-β treatment with reduced stemness and increased myofibroblast signatures (**Figure 8B**). Next, we examined the effect of *Klf4* depletion in AdvSca1-SM cells *in vitro*, by comparing AdvSca1-SM cells isolated from WT and *Klf4* KO cardiac tissue (**Supplemental Figure IXB**). Similar to the TGF-β-induced response in WT AdvSca1-SM cells, *Klf4* KO AdvSca1-SM cells exhibited reduced expression of stemness related genes under basal conditions (**Figure 8C**, top row). *Klf4* depletion resulted in up-regulation of some ECM genes (e.g. *Col1a1*, *Col4a1*, *Col4a2*), but also suppressed expression of others (e.g. *Postn* and *Col3a1*)(**Figure 8C**). Further we examined select genes that exhibited a blunted AngII-response in *Klf4* KO mice from the scRNA-seq data (Figure 6C). *Klf4* KO resulted in reduction in expression of *Tnc*, *Cxcl2,* and *Ccl19* (**Figure 8C**). Lastly, the expression of SMC contractile genes, including *Myh11*, *Cnn1*, and *Acta2,* were induced in *Klf4* KO cells compared to WT, likely due to de-repression of contractile gene promoters through KLF4 loss. Taken together, the data support the vital role of KLF4 in maintaining the phenotype of AdvSca1-SM cells.

**Figure 8.**
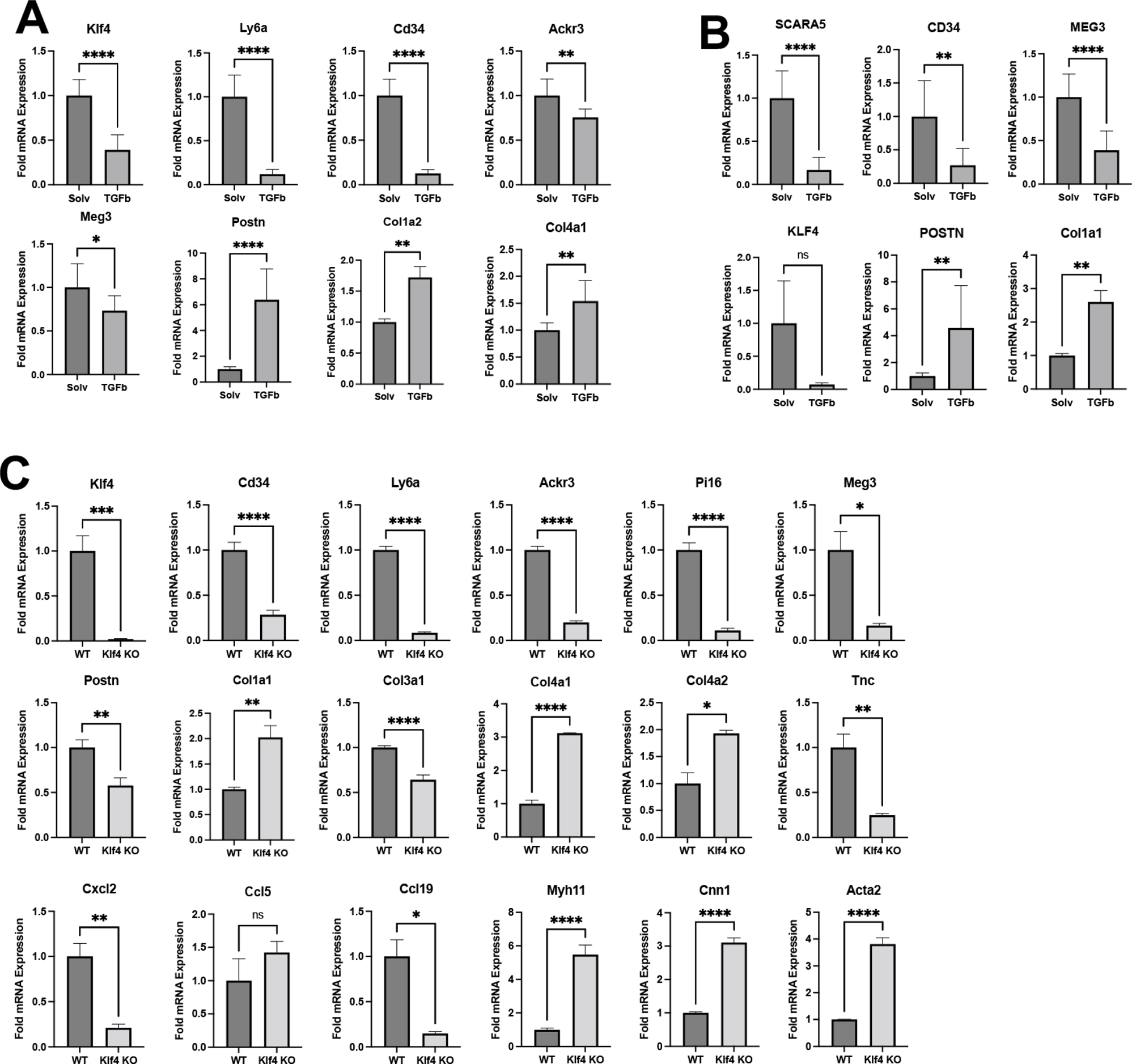
*Klf4* expression is associated with the stem-like and pro-inflammatory AdvSca1-SM phenotype *in vitro*. **(A)** AdvSca1-SM cell culture from Gli1Cre^ERT^-YFP mice was established as described in the Method section and subjected to TGF-β (5ng/mL) for 24 hours. qPCR was performed to examine select stemness and profibrotic gene expression. **(B)** Human AdvSca1-SM cell culture was established from donor coronary artery tissues. qPCR was used to determine the response to stemness and profibrotic gene to TGF-β. **(C)** WT and *Klf4* KO AdvSca1-SM cell culture were established from Gli1Cre^ERT^-YFP and Gli1-Cre^ERT^-YFP-Klf4^fl/fl^ mice. qPCR was performed to examine select stemness, profibrotic, proinflammatory and SMC contractile gene expression.

## DISCUSSION

Excessive cardiac fibrosis is the terminal manifestation of cardiac diseases and hampers the diastolic function of the heart. While there is little doubt that myofibroblasts are specialized cells for ECM production and play a central role in fibrotic diseases of the heart and other organs, the origin of myofibroblasts is still highly debated. Tamoxifen-inducible Cre-loxP based lineage tracing systems have been utilized in the study of ECM producing cells in the setting of fibrosis^42–44^. Kramann *et. al.* first reported that MSC-like *Gli1* expressing adventitial cells are important contributors to cardiac fibrosis^23^. Our recent publications also support the important function of adventitial *Gli1*^+^ cells in vascular injury-induced remodeling^21,22^ and atherosclerotic plaque composition^22^. In the current study, we used a scRNA-seq approach to define the transcriptomic changes in cardiac fibroblasts as well as *Gli1* AdvSca1-SM lineage cells. We identified a differentiation trajectory from AdvSca1-SM cells to myofibroblasts, characterized by the loss of expression of progenitor-associated genes and gain of expression of profibrotic genes. We found similar gene signature changes in cardiac and vascular tissues from cardiac hypertrophic patients, supporting the translational significance. Combining the lineage tracing approach with spatial transcriptomics methods, we further defined spatial embedding of AdvSca1-SM cells. AngII stimulated the expansion of AdvSca1-SM cells from the coronary artery perivascular region into the cardiac interstitium. The spatial transcriptomics data further support that AdvSca1-SM-derived myofibroblasts functionally interact with immune cells in promoting AngII-induced cardiac inflammation.

Previous studies of myofibroblast activation in the setting of fibrosis emphasized the gain of ECM gene expression, and expectedly, studies directed toward curbing the activation of TGF-β signaling and profibrotic gene expression has been the major focus of antifibrotic therapeutics^45–47^. However, our findings indicate a potential alternative therapeutic approach through maintenance of the stemness phenotype of AdvSca1-SM cells to thereby limit their contributions to the myofibroblast population. Importantly, stem-like AdvSca1-SM cells (marked by expression of *Pi16*, *Ly6a*, *Scara5*, *Ackr3*, *Meg3*, *Klf4*, etc.) have been identified in fibroblast populations across major organs^48^. Therefore, novel and clinically relevant anti-fibrotic therapeutics that target AdvSca1-SM cells to antagonize their profibrotic transition would serve as a universal anti-fibrotic treatment. As tissue/organ fibrosis accounts for ∼45% of all deaths in the Western world, this approach could have important clinical significance^49^.

Our scRNA-seq data, consistent with another report^50^, showed that *Gli1*^+^ AdvSca1-SM-derived cells contribute to almost all cardiac fibroblast clusters suggesting their major contribution to cardiac fibrosis. It is important to note, however, that the origin of cardiac myofibroblasts contributing to pathological cardiac fibrosis remains unclear. For instance, using lineage mapping systems, Kanisicak, et.al. described a periostin-positive myofibroblast population that was a major contributor to cardiac fibrosis in the setting of myocardial infarction (MI), AngII-mediated cardiac hypertrophy, and transverse aortic constriction (TAC)^37^. Using a novel and inducible periostin Cre driver, they reported that periostin lineage-traced cells accounted for the majority of myofibroblasts contributing to cardiac fibrosis. In addition, this group showed that the periostin-positive myofibroblasts were derived from resident cardiac fibroblasts of the TCF21 lineage, similar to our findings of AdvSca1-SM cells under both saline and AngII conditions. An important difference between their studies using the periostin-Cre system and our studies is the timing of tamoxifen treatment. We labeled *Gli1* lineage AdvSca1-SM cells prior to AngII treatment when AdvSca1-SM cells express a stemlike phenotype. Tamoxifen was allowed to wash out prior to AngII, which ensured that no additional cells were labeled during the course of the experiment even if *Gli1* was induced. In contrast, experiments in the Kanisicak, et.al. study induced stimulus (MI, AngII, TAC) prior to tamoxifen then continued the tamoxifen treatment throughout the experimental time course; therefore, any cell that induced periostin expression would be labeled for analysis. As periostin is induced in AdvSca1-SM cells within 2-3 days of disease stimulus as they differentiate into myofibroblasts^21^, it cannot be ruled out that a large percentage of the periostin lineage-derived myofibroblasts are, in fact, derived from activated *Gli1* lineage AdvSca1-SM cells. Nonetheless, further work is required to determine how much overlap between Gli1-lineage and periostin-lineage cells comprise the myofibroblast population in this AngII model.

Our initial report demonstrated that AdvSca1-SM cells originate from mature *Myh11-*expressing SMCs that are reprogrammed in late prenatal-to-early postnatal life as a normal physiological process^20^. We demonstrated this process using a highly specific SMC lineage system (inducible *Myh11*-Cre^ERT^) combined with ChIP analysis for a SMC-specific epigenetic lineage mark. Transcriptomic comparison of AdvSca1-SM cells from a SMC-specific lineage system to AdvSca1-SM cells from the *Gli1* lineage system used in the present study confirmed the SMC origin of AdvSca1-SM cells^21^. In contrast, using the same SMC *Myh11* lineage system, Kanisicak, et.al. found only a rare contribution of SMCs to myofibroblasts in the setting of MI. One likely explanation for the potential discrepancy is that *Myh11-*expressing SMCs would have to reprogram to AdvSca1-SM stem cells in response to MI and prior to differentiation into myofibroblasts in order to contribute to the reporter-positive myofibroblast population. Given the very short experimental timeline in this study (one week)^37^, this process is unlikely to contribute significantly to the myofibroblast population. Alternatively, SMC-to-AdvSca1-SM-to-myofibroblast differentiation is potentially not observed in the setting of MI.

It has been shown that genetic ablation of *Gli1*^+^ cells blunted pressure-overload-induced cardiac fibrosis in mouse models^23^. Compelling as the data were, systemic cell ablation can have off target effects and unintended consequences, and therefore unlikely to be viable as therapy in human fibrotic diseases. An improved understanding of the regulation of *Gli*1^+^ AdvSca1-SM cells phenotype is required to design better therapy. We previously reported that *Klf4* depletion in AdvSca1-SM cells induces a slow progressing spontaneous profibrotic phenotype in the absence of disease stimulus^21^. Quite surprisingly and paradoxically, in the current study, we found AdvSca1-SM-specific *Klf4* depletion conveys protection against cardiac fibrosis. The protection is associated with reduced proinflammatory interferon signaling and macrophage infiltration in the cardiac tissue. The defects in interferon signaling are not restricted to AdvSca1-SM cells where *Klf4* is depleted, but observed across multiple cell populations including macrophages, endothelial cells, SMCs, and pericytes. These findings strongly support the pivotal role of AdvSca1-SM cells in regulating and coordinating cardiac inflammation and indicate that manipulating the phenotype of AdvSca1-SM cells may have a profound effect on the phenotype and function of other cells in the microenvironment. It has been reported that global IFN-γ knockout mice are protected against AngII-induced recruitment of immune cells and cardiac fibrosis^51,52^. Cardiac fibroblasts readily respond to IFN-γ stimulation and elevate chemokine production^53^. However, the significance of IFN-γ signaling in cardiac fibroblasts and AdvSca1-SM cells is largely unexplored. The molecular function of AdvSca1-SM-specific *Klf4* in the regulation of interferon signaling through cell-to-cell communication warrants further investigations.

In addition to the inflammation changes, *Klf4* depletion also had a direct impact on mRNA expression of non-fibrillar collagen and matrix metallopeptidases (MMPs) in AdvSca1-SM cells. The increase of MMPs suggest active ECM breakdown in *Klf4* KO cardiac tissue. Currently, most imaging methods for ECM detection are specific to fibrillar collagen. To uncover changes in non-fibrillar collagen deposition, more sensitive methods, such as ECM Mass spectrometry, will be required. Further examination of the ECM deposition and the functional implications of the altered ECM production profile of AdvSca1-SM cells will be investigated in future studies. While the current study mainly focuses on AdvSca1-SM cells, the contribution of classical fibroblasts and myofibroblasts in cardiac fibrosis should not be disregarded. *Gli1*-lineage cells comprise a small percentage of the non-myocyte population, suggesting that they are unlikely to be the sole contributors to ECM deposition. The rarity of AdvSca1-SM cells stands in sharp contrast to the significant anti-fibrotic and anti-inflammatory phenotype observed upon AdvSca1-SM-specific *Klf4* depletion. One plausible explanation is that AdvSca1-SM cells, and their *Klf4* expression, coordinate the fibrotic response via paracrine signaling. Future studies will further investigate the cell-to-cell communication between AdvSca1-SM cells and other cardiac cell types, including fibroblasts.

In conclusion, our study sheds new light on the role of AdvSca1-SM cells in cardiac fibrosis and highlights the pivotal involvement of the transcription factor KLF4 in orchestrating their phenotypic transition and response to profibrotic stimuli. Using unbiased approaches, we uncovered a distinct differentiation trajectory of AdvSca1-SM cells under Angiotensin II-induced fibrotic conditions, characterized by the loss of stemness-related genes and acquisition of a profibrotic phenotype. Surprisingly, genetic knockout of Klf4 specifically in AdvSca1-SM cells prior to AngII treatment resulted in protection against cardiac inflammation and fibrosis, underscoring the critical role of KLF4 in promoting the profibrotic response of these cells. These findings not only deepen our understanding of the cellular mechanisms driving cardiac fibrosis but also highlight KLF4 as a potential therapeutic target for mitigating this pathological process. Remarkably, the observed gene program that controls AdvSca1-SM phenotype is conserved between murine and human, paving the way for future therapeutic discoveries targeting cardiac fibrosis. Our translational data from human cardiac hypertrophic tissue supports the clinical relevance of AdvSca1-SM cells in human cardiomyopathy. By delineating the molecular pathways governing AdvSca1-SM cell behavior in response to fibrotic stimuli, our work opens new avenues for developing targeted therapies aimed at modulating these cells to prevent or reverse cardiac fibrosis. Moreover, the presence of perivascular AdvSca1-SM cells in multiple organs suggests that targeted anti-fibrotic agents developed based on these findings could have applications beyond the cardiovascular system. Ultimately, further investigation into the intricate regulatory networks involving KLF4 and AdvSca1-SM cells promises to uncover innovative strategies for combating cardiac fibrosis and improving patient outcomes.

## AUTHOR CONTRIBUTIONS

SL, MWM, and MCMWE designed the studies. SL, AJJ, KAS, and MFM performed the experiments. TN assisted with mouse colonies and molecular biology experiments. SL performed the bioinformatics analysis. KSM provided human tissues for human studies and assisted with the scRNA-Seq. SL and MCMWE wrote the manuscript. AJJ, AMD, KAS, MFM, TN, RAN, KSM, and MWM edited the manuscript.

## ACKNOWLEDGMENTS

University of Colorado Shared Resources (SR) are supported by the National Cancer Institute through the Cancer Center Support Grant (P30CA06934). The Genotype-Tissue Expression (GTEx) Project was supported by the Common Fund of the Office of the Director of the National Institutes of Health, and by NCI, NHGRI, NHLBI, NIDA, NIMH, and NINDS. The data used for the analyses described in this manuscript were obtained from the GTEx Portal (V8) on 04/06/2021.

## SOURCES OF FUNDING

This work was supported by grants R01 HL121877 (Weiser-Evans, Majesky), R01 HL151331 (Weiser-Evans), R01 HL148167 (Kovacic; Weiser-Evans subcontract), a CFReT pilot award (Weiser-Evans), and the Loie Power Robinson Stem Cell & Regenerative Medicine Fund, Seattle Children’s Research Institute (Majesky).

## DISCLOSURES

None

## SUPPLEMENTAL TABLES

**Supplemental Table 1:** qPCR primer sequences.

**Supplemental Table 2:** DGE analysis comparing AngII-treated and saline-treated SM-Fib cluster cells. Positive log2FoldChange indicated up-regulation by AngII.

**Supplemental Table 3:** Up-regulated pathways in AngII-treated SM-Fib cluster cells.

**Supplemental Table 4:** DGE analysis comparing AngII-treated and saline-treated Macrophages and monocytes clusters cells. Positive log2FoldChange indicated up-regulation by AngII.

**Supplemental Table 5:** Up-regulated pathways in AngII-treated Macrophages and monocytes clusters cells.

**Supplemental Table 6:** DGE analysis comparing AngII-treated and saline-treated YFP^+^ cells. Positive log2FoldChange indicated up-regulation by AngII.

**Supplemental Table 7:** Up-regulated pathways in AngII-treated YFP^+^ cells.

**Supplemental Table 8:** Down-regulated pathways in AngII-treated YFP^+^ cells.

**Supplemental Table 9:** DGE analysis comparing YFP^+^ cells in MyoFib cluster and SM clusters. Positive log2FoldChange indicated up-regulation in MyoFib.

**Supplemental Table 10:** Up-regulated pathways in YFP^+^ cells in MyoFib cluster versus SM clusters.

**Supplemental Table 11:** Down-regulated pathways in YFP^+^ cells in MyoFib cluster versus SM clusters.

**Supplemental Table 12:** DGE analysis comparing hypertrophic and non-hypertrophic cardiac samples from GTEx data. Positive log2FoldChange indicated up-regulation in hypertrophic tissues.

**Supplemental Table 13:** DGE analysis comparing AngII-treated YFP^+^ SM-Fib cells from *Klf4* KO and WT mice. Positive log2FoldChange indicated up-regulation in *Klf4* KO tissues.

**Supplemental Table 14:** Up-regulated pathways in AngII-treated YFP^+^ SM-Fib cells from *Klf4* KO compared to WT.

**Supplemental Table 15:** Down-regulated pathways in AngII-treated YFP^+^ SM-Fib cells from *Klf4* KO compared to WT.

**Supplemental Table 16:** DGE analysis comparing AngII-treated Mac-Mono cells from *Klf4* KO and WT mice. Positive log2FoldChange indicated up-regulation in *Klf4* KO tissues.

**Supplemental Table 17:** Down-regulated pathways in AngII-treated Mac-Mono cells from *Klf4* KO compared to WT.

**Supplemental Table 18:** Up-regulated pathways in AngII-treated Mac-Mono cells from *Klf4* KO compared to WT.

**Supplemental Table 19:** Down-regulated pathways in AngII-treated endothelial cells from *Klf4* KO compared to WT.

**Supplemental Table 20:** Down-regulated pathways in AngII-treated smooth muscle cells/pericytes from *Klf4* KO compared to WT.

## Notes

### Competing Interest Statement

The authors have declared no competing interest.

